# Transcriptional profiling provides insights on sublethal thermal stress thresholds in juvenile bull trout

**DOI:** 10.1101/2024.10.22.619659

**Authors:** C. Best, T. C. Durhack, A. J. Chapelsky, M. Aminot, S. S. Islam, D. D. Heath, N. J. Mochnacz, K. M. Jeffries

## Abstract

Bull trout (*Salvelinus confluentus*) inhabit mainly cold-water streams in their Western North American range. Multiple anthropogenic stressors, including climate change, have caused population declines and range reduction, leading to classification of bull trout as threatened. Characterizing how fish respond to increasing temperature can identify key thresholds beyond which fish health is impacted, which is useful for developing species recovery strategies.

Juvenile bull trout were acclimated to a range of relevant temperatures (6-21 °C), after which mRNA transcripts involved in responding to thermal stress were measured using high-throughput qPCR, and their upper thermal tolerance (critical thermal maximum, *CT_max_*) was determined. Beyond 18 °C appears to be a critical sublethal threshold for bull trout: capacity to increase *CT_max_* through acclimation was lost, growth and survival decreased, and transcriptional data suggest widespread activation of cellular stress and growth suppression. Below 18 °C, transcriptional data suggest subtler metabolic adjustments in response to acclimation temperature, preceding changes in whole-animal performance. Altogether, these data provide insight into thermal responses in bull trout, and their capacity to cope with climate-change related warming.

## 1 Introduction

Fishes encounter many challenges in their environment and warming climates are of particular concern, with potential and realized impacts on their physiology, life history, habitat, and ultimately population ranges (Harrod 2016; Cooke et al. 2022). Characterization of the physiological responses of fishes to elevated temperatures in a controlled environment provides important information about how they may respond to a warming natural environment. Fish respond to increasing temperatures in a myriad of ways, with effects at many levels of biological organisation. Thermal performance curves are a common way to illustrate the effect of temperature on animal performance (typically through effects on growth, metabolism, or enzyme activity), allowing identification of temperatures that optimize or limit performance (Schulte et al. 2011). Underlying these changes in performance are cellular and molecular level responses: changes in signalling pathways (e.g. the heat shock response) that are recruited at different temperature thresholds (Jeffries et al. 2018). These thermal response thresholds can identify the induction of stress responses, and mark when temperature increases begin to adversely impact the animal and may threaten survival (Jeffries et al. 2018). While these thresholds were originally applied to an acute thermal response, they can reflect impacts across chronic exposures (Mackey et al. 2021). On a chronic time scale, the response of fishes to elevated temperatures often includes acclimation - a form of phenotypic plasticity during which an animal adjusts their biological systems in response to an environmental change to optimize their health and performance. Thermal acclimation is a dynamic, adaptive process and can dictate the shape of thermal performance curves (Schulte et al. 2011). The temperature at which acclimation ceases to occur is an important sublethal threshold and is associated with chronic thermal stress and potential reductions in fitness (Komoroske et al. 2015; Morrison et al. 2020), observable from the molecular to the whole-animal level.

Transcriptional changes provide insight into the regulation of many physiological pathways that may be impacted by temperature, and are most useful when assessed in the context of other whole-body physiological performance metrics (Jeffries et al. 2018). Transcriptional changes may reveal more subtle physiological adjustments preceding the manifestation of changes in performance. Changes in mRNA transcript abundance can offer insight into mechanisms of thermal acclimation, induction of stress responses, and whether fish are experiencing acute or chronic stress. The sensitivity and breadth of responses that can be addressed by transcript abundance, combined with the increasing accessibility of measuring many target genes, makes transcriptional evaluation of thermal responses a powerful approach for assessing thermal response thresholds, particularly in combination with whole-animal performance traits (Komoroske et al. 2015; Jeffries et al. 2018, 2021; Teffer et al. 2019; Mackey et al. 2021; Bouyoucos et al. 2023; Durhack et al. 2024). Transcriptional profiling has also been suggested to be a rapid method to characterize thermal stress responses in species with limited information available, or those which are difficult to study in a laboratory setting (Jeffries et al. 2016). Transcriptional profiling in the current study was conducted using a stress-response transcriptional profiling (STP) chip, which targets genes that indicate fish health and stress, and is designed to work across salmonids (Islam et al. 2024). The STP-chip was developed as part of the genomic network for fish identification, stress, and health (GEN-FISH; https://gen-fish.ca/) project which aims to assess performance of Canada’s freshwater fish species as they deal with increasing stressor burdens.

Cold-water fishes are more susceptible to adverse effects of climate warming because they occupy cold, and often narrow, thermal niches (Cooke et al. 2022). Bull trout (*Salvelinus confluentus*) are a stenothermal char native to Western North America that typically inhabits cold streams that are thermally stable (Mochnacz et al. 2023). Populations of bull trout have declined and were first listed as threatened in the United States in 1998 (USFWS 1998) and in Canada in 2012 (COSEWIC 2012). Climate warming is thought to be an important factor in this decline, particularly in their southern range (Rieman et al. 2007; Kovach et al. 2017; Mochnacz et al. 2023). Of the North American salmonids, bull trout occupy one of the coldest thermal niches (Mochnacz et al. 2023). They are most often found in streams with mean August temperatures of 6 to 8 °C and are rarely found above 11 °C (Isaak et al. 2015; Mochnacz et al. 2023). While they have been observed at temperatures as high as 20.5 °C in the summer (Adams and Bjornn 1997), this is not typical nor optimal for bull trout, as their ultimate upper incipient lethal temperature (UUILT, 50% survival) has been predicted to be 20.9 °C (Selong et al. 2001). Bull trout appear to have among the lowest thermal limits of the salmonids, alongside Arctic char (*Salvelinus alpinus*) and Arctic grayling (*Thymallus arcticus*) in terms of both chronic (UUILT) and acute (critical thermal maximum, *CT_max_*) metrics (Selong et al. 2001). Much of the work to date on bull trout focuses on field studies, while controlled lab studies characterizing their thermal performance are limited, due mainly to their status as a threatened species in much of their natural range (Selong et al. 2001; McMahon et al. 2007; Mesa et al. 2013; Durhack 2020).

Additionally, their transcriptional responses to temperature have not been characterized. Indeed, in comparison to other salmonids, relatively little is known about the physiological and molecular responses to thermal stress or other environmental stressors in bull trout, despite their status as a threatened species.

The aim of this study was to characterize the transcriptional profile of bull trout as it changes with increasing acclimation temperature, then integrate this information with other performance endpoints to identify thermal thresholds for this species. To test this, we investigated the response of juvenile (young-of-the-year) bull trout (reared at 10 ± 1 °C) acclimated to a range of ecologically relevant temperatures (6, 9, 12, 15, 18, 21 °C) by measuring transcriptional changes across three tissues (i.e., gill, liver and muscle).

Transcriptional changes were then compared to whole-body physiological performance (growth, survival) and thermal limits (*CT_max_* and agitation temperature i.e., the temperature at which fish begin to elicit an escape response to an acute thermal stress; (McDonnell and Chapman 2015)). The range of acclimation temperatures chosen included water temperatures where this species is typically found (6 - 12 °C), as well as higher temperatures that may become relevant with climate warming or are already experienced by bull trout at the southern end of their distribution. We predicted that bull trout would perform best at temperatures where they typically occur in the wild, demonstrate activation of adaptive thermal acclimation at intermediate temperatures, then reach a sublethal threshold at the highest temperatures at which they lose their capacity for acclimation and experience chronic thermal stress. The results of this study will inform management of bull trout populations in the wild, particularly to conserve imperilled populations.

## 2 Materials and methods

### 2.1 Animal Husbandry

Young-of-the-year (YOY) bull trout used in the experiment were the F1 offspring of gametes collected September 15-16, 2021 from wild bull trout in Smith-Dorrien Creek, Alberta, Canada. Gametes were shipped back to the Freshwater Institute in Winnipeg, Manitoba, Canada, where eggs were fertilized and then put in incubation trays. Embryos were incubated at ∼4 °C until hatching started (January 11, 2022), were warmed to 5 °C during hatching, and held there until February 23, 2022 when temperatures were warmed at a rate of 1.0 °C/day up to 8 °C. Alevins were kept in hatching trays until March 15, 2022 when they were transferred to tanks for swim up in April – May, 2022. Following swim up, fish were transferred to general fish holding tanks where the water temperature was gradually increased to 10 ± 1 °C for long-term holding in 100 L tanks maintained on a flow-through of de-chlorinated water and with independent aeration (Kissinger et al., 2024 *in prep*). Fish were maintained under a 12:12 h photoperiod (65 min of dawn/dusk, full light starting at 0705 hours, and full dark at 1905 hours) and fed daily at a rate of 0.5% body mass of commercial food pellets (#2 crumble EWOS Pacific: Complete Fish Feed for Salmonids, Cargill, Winnipeg, Manitoba, Canada).

### 2.2 Experimental Protocols

#### 2.2.1 Acclimation

Experimental fish were acclimated in 200 L tanks for each experimental treatment. Fish were haphazardly netted from the general population tanks and weighed and measured before being transferred into their acclimation tanks. Total tank mass was used to determine the appropriate amount of feed to achieve a 2% body mass feeding rate. Acclimation tanks were held at the same temperature as the general population overnight while fish acclimated to the new tank. Tanks were then adjusted at a rate of 2 °C per day until they reached the treatment temperature using a TempReg instrument (Loligo^®^ Systems, Tjele Denmark) connected to coils in external heating and cooling tanks. Fish were acclimated to treatment temperatures for just over 3 weeks before experiments began (24 days, except 21 °C was 22 d, 9 °C was 23 d). All procedures were conducted in accordance with the guidelines established by the Fisheries and Oceans Canada Ontario & Prairies Animal Care Committee (OPA-ACC AUP-2022-58) following the standards set by the Canadian Council for Animal Care.

#### 2.2.2 Study design

From June 8 – July 11, 2022, six treatments (6, 9, 12, 15, 18, 21 °C) of juvenile bull trout (*n* = 35 fish per treatment) were used to assess the effects of different thermal acclimations (total of *n* = 210). Sampling in each treatment was separated into two experiments: Transcriptional profiling – where *n* = 15 fish were sampled immediately from each treatment tank following the end of acclimation, or *CT_max_* – where *n* = 15 fish were put through a critical thermal maximum testing protocol (Beitinger et al. 2000). Five extra fish were acclimated in each tank in case of mortalities during acclimation, which did occur resulting in a reduction in the planned *n* = 15 for some groups (see Table 2). At the end of each time point, fish were euthanized in tricaine methanesulfonate (MS-222; concentration: 450 mg L^-1^ buffered with 900 mg L^-1^ sodium bicarbonate) for three minutes and length (fork and total in mm) and mass (g) were recorded, and tissues were taken for transcriptional profiling.

**Table 1.**
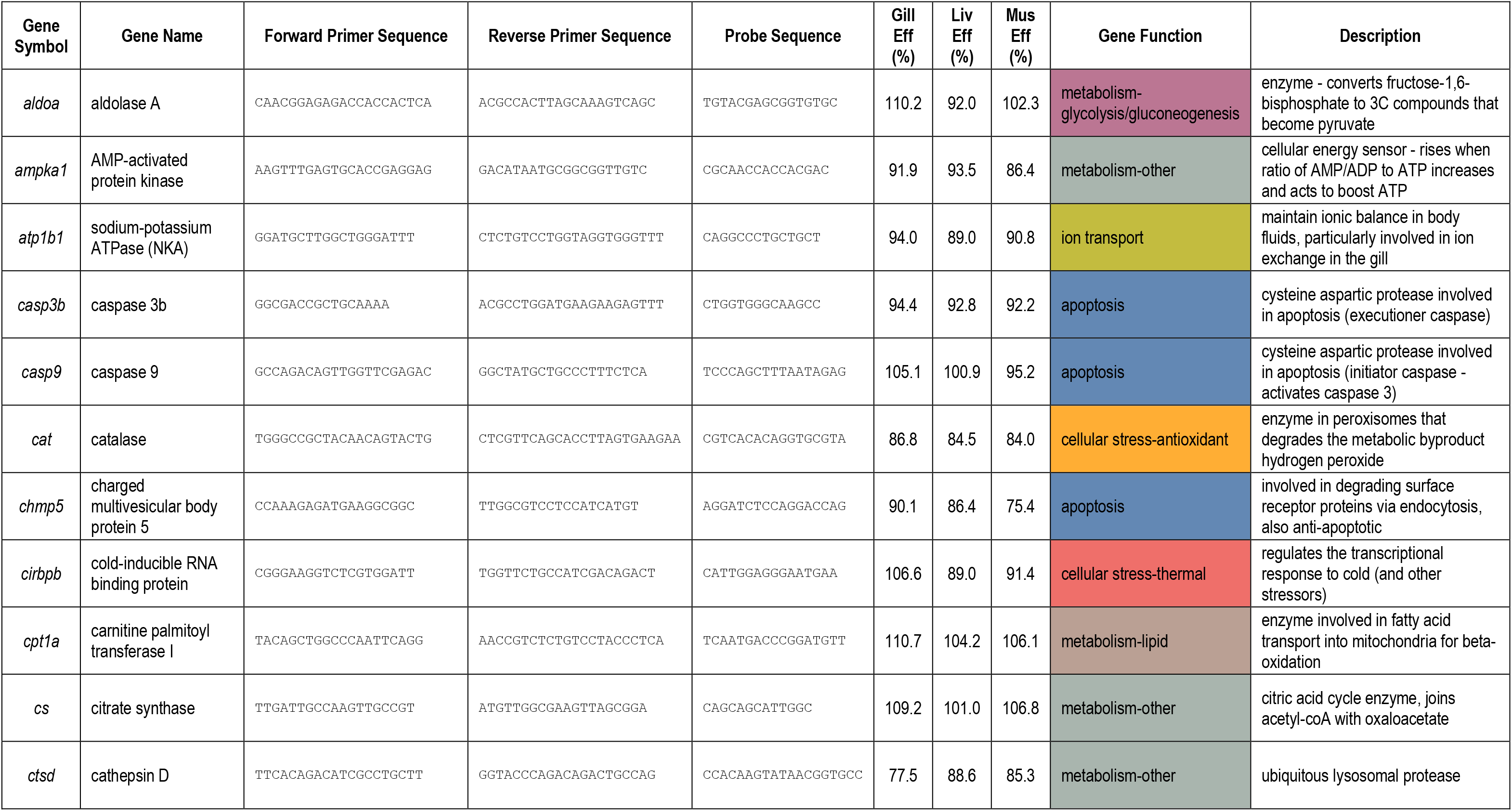
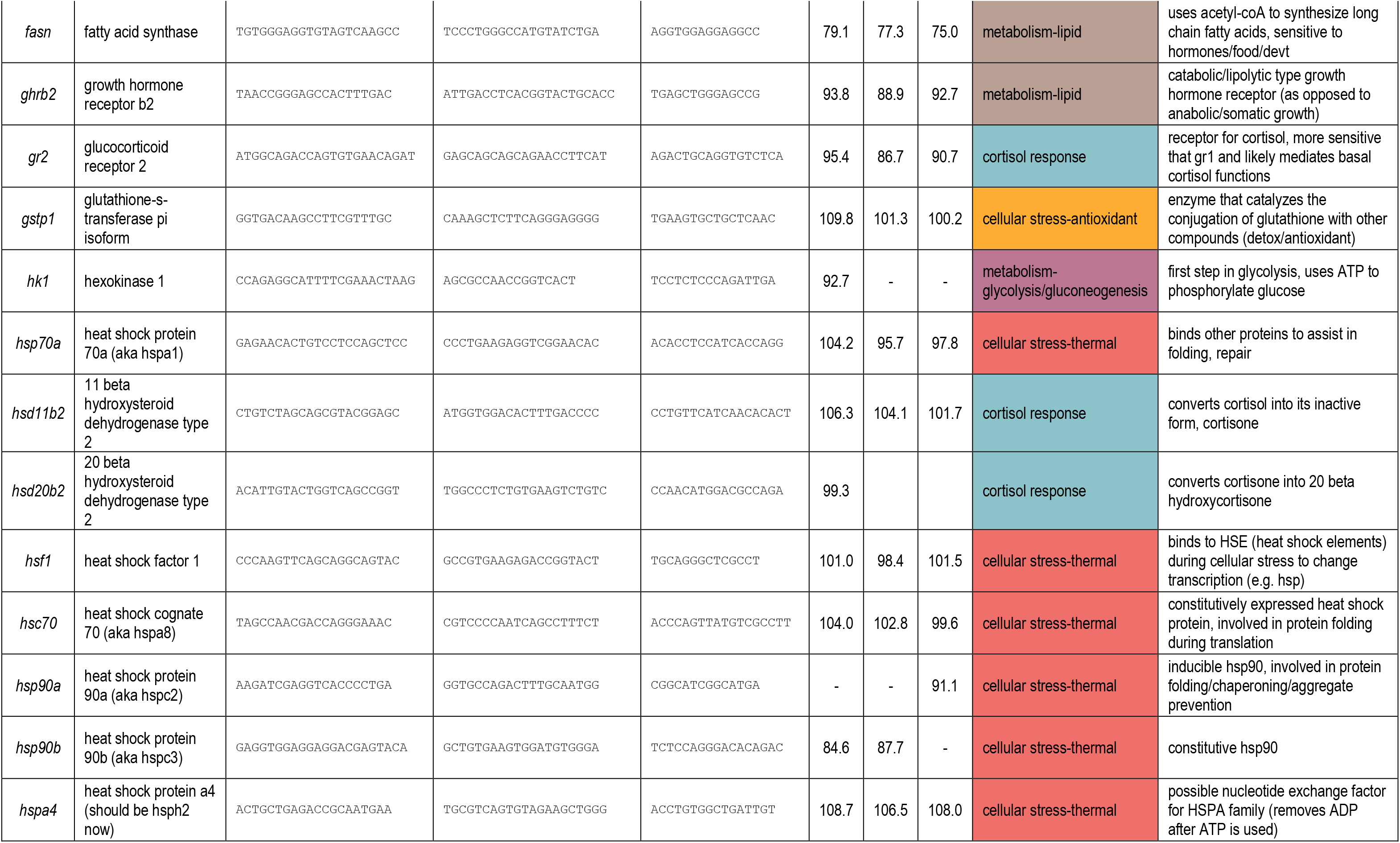
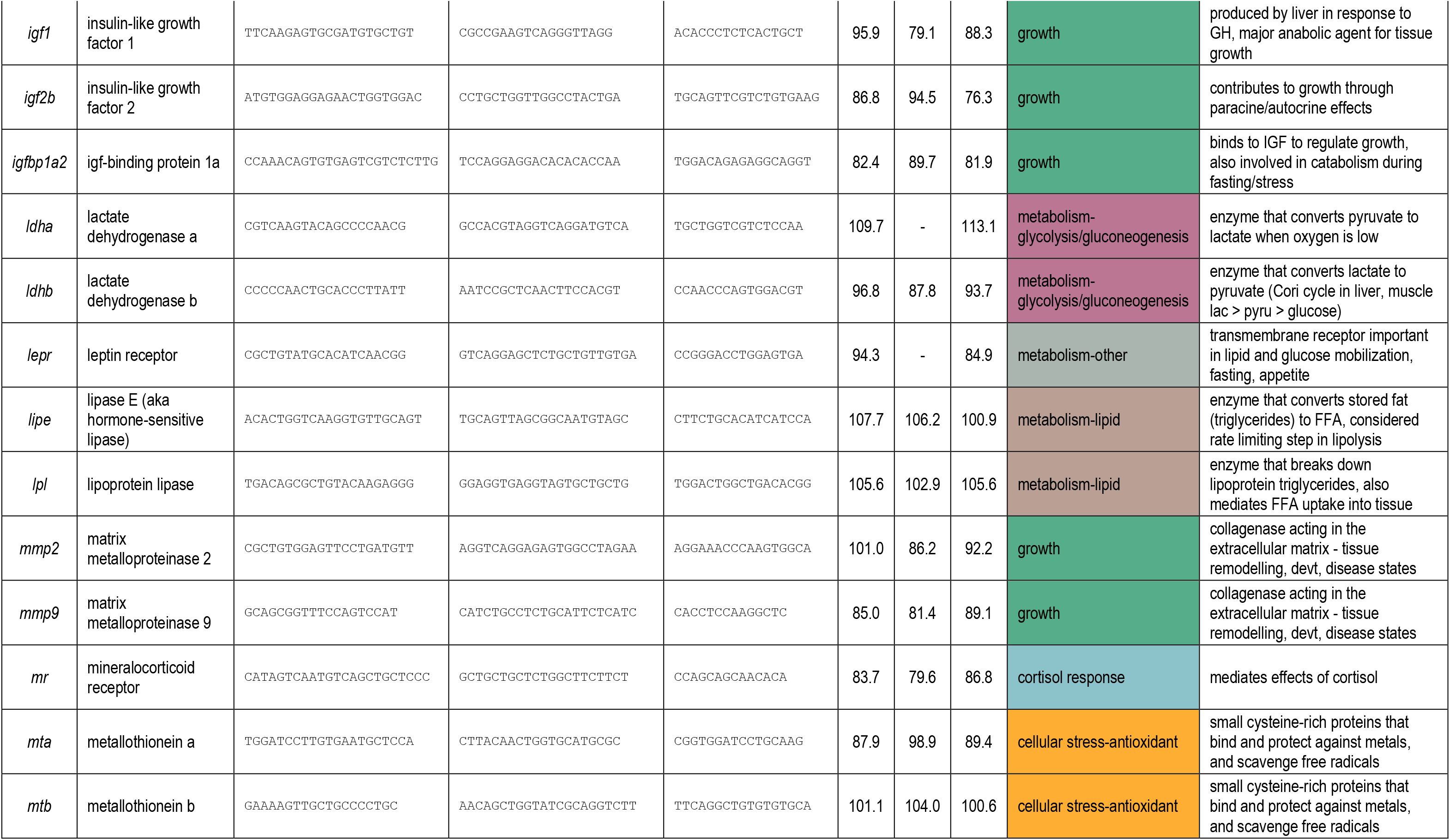
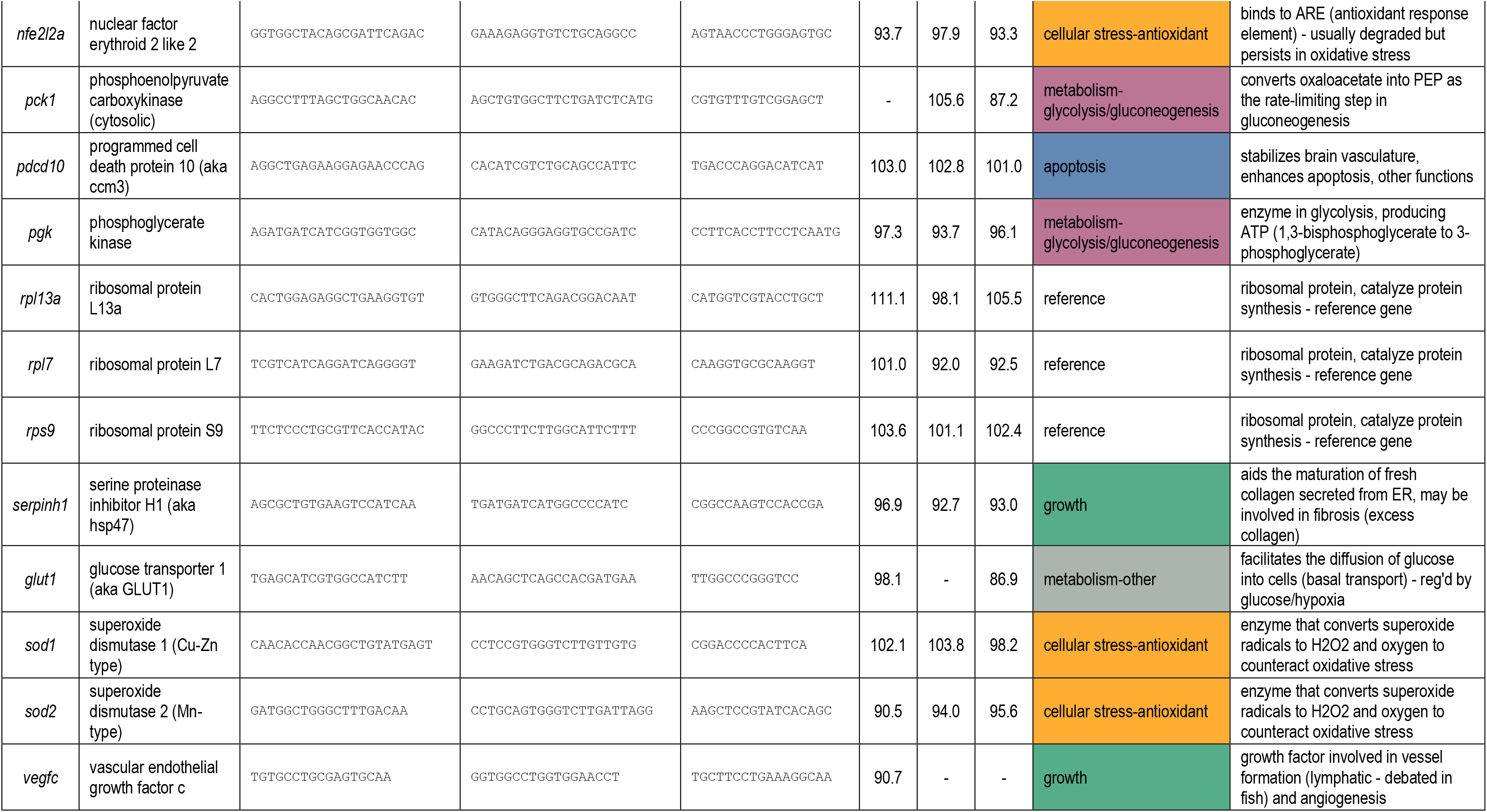
Primers and probe sequences used for OpenArray qPCR reactions.

**Table 2.** Weight and mortality data over the acclimation period for YOY bull trout (*Salvelinus confluentus*) exposed to a range of acclimation temperatures. Starting group size was *n* = 35 for all groups. Acclimation began after completion of the ramping period (2 °C/day) from initial holding temperature (10 ± 1 °C), and end date is when fish were sampled for molecular analysis (*n* = 15), with *CT_max_* trials performed the following day (*n* = 15, *n* = 13 for 18 °C and *n* = 9 for 21 °C).

| Group | Acclim.<br>Start Date | Sampling<br>Date | Days at<br>Acclim. | Growth<br>Days | Start Weight<br>(g, mean±SE) | End Weight<br>(g, mean±SE) | No.<br>Deaths | Mortality<br>(%) |
| --- | --- | --- | --- | --- | --- | --- | --- | --- |
| 6 °C | 14-Jun-22 | 8-Jul-22 | 24 | 28 | 0.42 ± 0.03 | 0.66 ± 0.03 | 0 | 0.0 |
| 9 °C | 16-Jun-22 | 9-Jul-22 | 23 | 23 | 0.48 ± 0.03 | 0.69 ± 0.04 | 2 | 5.7 |
| 12 °C | 14-Jun-22 | 8-Jul-22 | 24 | 25 | 0.48 ± 0.03 | 0.78 ± 0.05 | 1 | 2.9 |
| 15 °C | 16-Jun-22 | 10-Jul-22 | 24 | 27 | 0.53 ± 0.03 | 0.82 ± 0.05 | 1 | 2.9 |
| 18 °C | 15-Jun-22 | 9-Jul-22 | 24 | 29 | 0.49 ± 0.03 | 0.69 ± 0.04 | 7 | 20.0 |
| 21 °C | 15-Jun-22 | 7-Jul-22 | 22 | 28 | 0.54 ± 0.03 | 0.51 ± 0.03 | 9 | 25.7 |

#### 2.2.3 CT_max_ experiments

Following thermal acclimation, *n* = 15 fish per treatment were put through a *CT_max_* testing protocol (Beitinger et al. 2000). Fish were tested in groups of fifteen in 200 L tanks, where each fish was placed in an individual acrylic chamber (volume – 212 ml) with netting over the ends to contain them and allow for tracking of individual fish behaviour. *CT_max_* fish were placed in their sampling chambers to habituate to the chamber overnight prior to experimental trials (approx. 18 hours). Temperature of the tank was recorded using a HOBO Tidbit® v2 temperature logger (ONSET Computer Corporation, Bourne, Massachusetts, USA). The next morning, tank water level was reduced to 11 cm depth to facilitate proper heating. *CT_max_* experiments were conducted with a heating rate of ∼0.3 °C min^-1^ using three 300 W titanium heaters (Finnex TH-0300S titanium heaters, Finnex, Chicago, USA). Experimental tanks had air stones to maintain dissolved oxygen levels and two 5 L·min^-1^ pumps (Eheim, Deizisau, Germany) circulating water to ensure homogenous temperature throughout the tank. During heating, fish were monitored for two behavioural reactions – agitation temperature and *CT_max_*. Agitation temperature was determined to be reached when the fish displayed sustained (>5 s) active movement (burst swimming or repeated turning around in the chamber). The *CT_max_* was defined as when the fish was unable to maintain equilibrium (loss of equilibrium, LOE). Fish were removed from the treatment immediately upon reaching *CT_max_* and placed in a recovery bath at 10 °C for one hour where all but one fish recovered (upright swimming behaviour and normal reaction to visual stimuli) within 30-45 minutes. Recovered fish were then euthanized and measured and weighed, as described above. Oxygen saturation was measured at the beginning and end of each trial and never fell below 90%.

#### 2.2.4 Transcriptional profiling

Following thermal acclimation, *n* = 15 fish per treatment were haphazardly netted from the acclimation tanks, euthanized in a buffered MS-222 bath, and tissue samples were taken for transcriptional profiling work. Gill and liver tissue was sampled by taking the entire organ from the fish and immediately stored in a pre-weighed vial of RNA*later*^TM^ (Invitrogen™, Carlsbad, California, USA). For muscle tissue, the front half of the fish muscle (back of head to dorsal fin) was removed and placed in RNA*later*^TM^. Tissue samples in RNA*later*^TM^ were kept stored at 4 °C for ∼24 h following sampling prior to storage in a -80 °C freezer.

### 2.3 Measurement of transcript abundance

Tissues were processed for RNA extraction using a KingFisher Duo Prime instrument and MagMAX mirVana Total RNA Isolation Kits (Applied Biosystems) following manufacturer’s protocols. Total RNA quantity and quality were assessed through NanoDrop and gel electrophoresis. Following manufacturer protocols, RNA was treated with DNase to remove genomic DNA (ezDNase, Invitrogen) and then 1 µg was converted to cDNA (High Capacity cDNA Reverse Transcription Kit, Applied Biosystems) and stored at -20 until PCR analysis.

Taqman qPCR assays were designed against 56 genes based on homology to multiple salmonid species, specifically those for which mRNA sequences were available in GenBank or through publicly available reference transcriptomes. Target genes were involved in thermal stress and general fish health and included four reference genes (see Table 1). Custom OpenArray high-throughput qPCR chips (Thermo Fisher Scientific) were printed, such that the “wells” (i.e., through-holes) came pre-coated with primers and probes (see Table 1 for sequences). Each OpenArray plate accommodated the amplification of all 56 genes in up to 48 samples, such that each plate could simultaneously amplify 2688 Taqman qPCR reactions. Samples were run in duplicate (i.e. 24 samples in duplicate per plate), by first assembling 5 µl reactions consisting of 2.5 µl 2X OpenArray master mix (Thermo Fisher Scientific) and 2.5 µl 5X diluted cDNA, in a 384-well plate. These reactions were then automatically spotted onto the custom OpenArray plate by the OpenArray Accufill System (Thermo Fisher Scientific), such that each 5 µl reaction corresponded to one sub-array containing all 56 assays. Within 2 minutes of loading (to avoid evaporation) the lid was affixed, and the plate was filled with oil and sealed following manufacturer’s protocols. OpenArray plates were then run on the QuantStudio 12K Flex System (Thermo Fisher Scientific). Post-run images were analyzed to confirm proper array loading (ROX fluorescence for filling of through-holes and light images for detection of any leaks/anomalies). Through-holes that were not loaded properly were omitted from analysis, and no leaks or anomalies were observed.

### 2.4 Data Analysis

Statistical analyses were performed in R v4.2.3 (R Core Team 2023), RStudio v2023.6.0.421 (Posit Team 2023), and Sigmaplot v13.0 (Systat Software, Inc.) and a significance level (α) of 0.05 was used. For *CT_max_* and agitation temperature, data did not meet the assumptions of a parametric test and therefore a nonparametric Kruskal-Wallis test followed by Dunn’s post hoc test was performed to determine significant differences between acclimation temperature groups. In the subset of *CT_max_* data without the 21 °C group, parametric assumptions were met and a one-way ANOVA was performed, followed by a Tukey post hoc test. Relative growth rates (RGR, %) were calculated as RGR (%) = [(Y2 - Y1)/(Y1 x t)] x 100, where Y1 and Y2 are initial and final mean individual fish masses (g) and t is number of days between mass measurements for each tank (Bear et al. 2007). The relative growth curve across acclimation temperature was fit with a quadratic regression.

Transcript abundance data were analysed for each of the three tissues (gill, liver, muscle). Using QuantStudio Design and Analysis v1.5.2 and ExpressionSuite v1.3 software (Thermo Fisher Scientific), raw amplification data (Crt values, relative cycle thresholds) were screened for quality, including removal of any samples where through-holes were not filled, or where amplification scores (reflecting amplification curve quality) and Cq confidence scores (reflecting reliability of the Crt value) were below acceptable limits as determined by the manufacturer’s protocols (1.24 and 0.8, respectively). Duplicate values with high variability were also omitted from analyses. Genes where over half of total samples were removed based on these quality parameters were removed entirely from further analyses.

Following quality screening, Crt values were analysed in R using the mcmc.qpcr package (Matz et al. 2013). LinRegPCR software was used to calculate amplification efficiencies based on the slope of the linear portion of the amplification curve using baseline-corrected fluorescence data as recommended for QuantStudio chip-based qPCR (Ruijter et al. 2009). These amplification efficiencies were accounted for in the mcmc.qpcr analysis in R, and are reported in Table 1. All genes for a given tissue were processed simultaneously by Bayesian model-based analysis according to the recommended workflow of mcmc.qpcr. First, a naïve model without reference gene normalization was generated, and given the absence of global effects, this was followed by a more powerful informed model that included reference gene normalization (ribosomal proteins *rpl13a*, *rpl7*, *rps9*) which comprised the final analysis. R scripts and data for this analysis are on GitHub (https://github.com/c3best/Bull_Trout_Transcript_Analysis), and geneWise output matrices for each tissue containing pairwise fold-differences and MCMC-based *p*-values indicating significant differences between acclimation temperature groups are in the supplementary information (Tables S1, S2, S3 for gill, liver, and muscle respectively).

Transcript abundance data were further analysed using a principal components analysis (PCA) in R using the ‘FactoMineR’ package (Lê et al. 2008) to determine if the variation in the data was sufficient to separate out groups by acclimation temperature, and if so, which transcripts were most involved (highest loadings). The delta-Crt values were used, i.e. Crt values corrected by the geometric mean of the 3 reference genes. The R scripts for this analysis are on GitHub (https://github.com/c3best/Bull_Trout_Transcript_Analysis).

## 3 Results

### 3.1 Thermal Tolerance

As acclimation temperature increased, *CT_max_* also increased in a generally step-wise fashion (Fig. 1A). At the lowest acclimation temperature (6 °C) the average *CT_max_* (± SEM) was also lowest, at 25.5 ± 0.1 °C. Mean *CT_max_* continued to increase by 0.7-1.3 °C for every 3 °C increase in acclimation temperature until the 18 °C acclimation temperature, at which *CT_max_* plateaued (29.3 ± 0.1 °C). Individuals in the 21 °C acclimation group had similar or lower *CT_max_* relative to 18 °C, as well as increased variability (SEM = 0.42 °C). This variability contributed to a reduced ability to detect differences statistically, and removal of this group from the analysis resulted in all remaining acclimation groups being significantly different from one another (one-way ANOVA with Tukey post hoc, *p* = <0.001).

**Figure 1.**
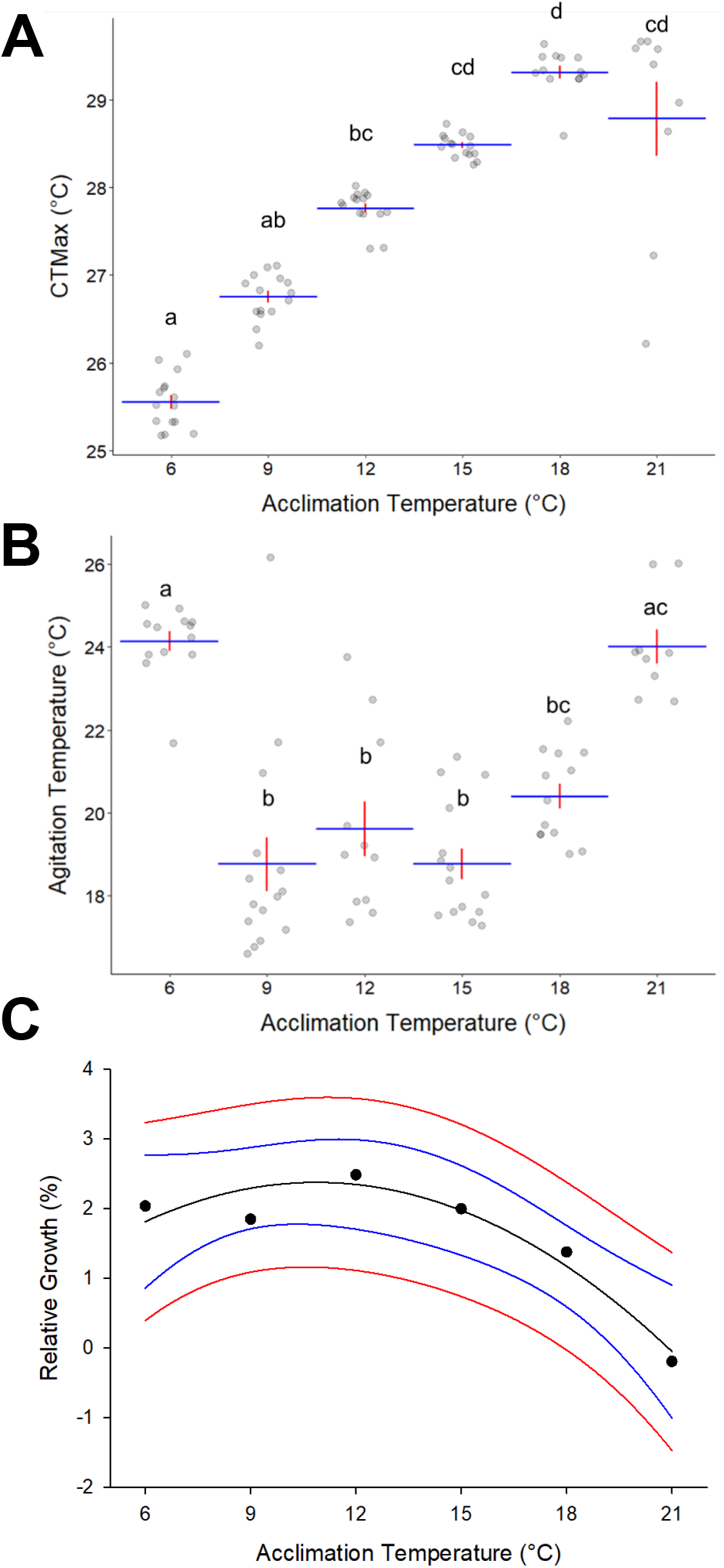
Critical thermal maximum (A), agitation temperature (B), and relative growth rates (C) of bull trout (*Salvelinus confluentus*) acclimated to temperatures ranging from 6 to 21 °C for three weeks. In A and B, horizontal and vertical lines indicate group means and standard error, respectively. Points represent values for individual fish (*n* = 9-15). Groups with different lowercase letters are significantly different from one another (Kruskal-Wallis test with Dunn’s post-hoc, *p* = <0.001 for both). In C, points represent the relative growth rates calculated using the mean weight of bull trout at the start and end of the experiment at each acclimation temperature group. Blue and red lines represent the 95% confidence and prediction bands, respectively. All fish were included in this analysis (*n* = 35 initial, *n* = 26-30 final). Growth was modelled with a quadratic curve (y = -0.0237x^2^ + 0.5155x – 0.4247, vertex = 10.9 °C, R^2^ = 0.8773, *p* = 0.02).

Agitation temperature displayed a U-shaped pattern, with fish in the coldest (6 °C) and warmest (21 °C) acclimation groups displaying agitation behaviour at the warmest temperatures (∼24 °C, Fig 1B). In comparison, fish acclimated at 9, 12, or 15 °C displayed agitation behaviour at significantly cooler temperatures (∼19 °C). In combination with *CT_max_*, this means that the 6 °C group had the smallest thermal agitation window, with agitation preceding *CT_max_* by just 1.4 °C. In comparison, this window averaged 8.0-9.7 °C in all other groups except 21 °C (4.8 ± 0.7 °C).

### 3.2 Growth and Survival

Relative growth rate was highest at 12 °C and lowest at 21 °C (Fig. 1C). The quadratic regression of relative growth rate vs. acclimation temperature was significant (R^2^ = 0.8773, *p* = 0.02), with estimated relative growth rate peaking at 10.9 °C (Fig. 1C). Mortality remained low throughout the experiment at the four lowest acclimation temperatures (0-2 mortalities per tank of *n* = 35, Table 2) and was highest at the two warmest acclimation temperatures: 20 and 25.7% in the 18 and 20 °C acclimation groups, respectively.

### 3.3 Transcript Abundance

In the gill, the transcript abundances of a total of 44 target genes were measured at a sufficient quantity and quality for analysis. When transcript abundance was assessed by PCA (Fig. 2A), 30.0% of the variation was explained by PC1 while 18.7% of the variation was explained by PC2. Across PC1, 6 and 9 °C were similarly grouped, as were 15, 18, and 21 °C. These groupings were distinct across PC1, with 12 °C being intermediate. The largest loadings on PC1 in the gill included transcripts involved in metabolism and cellular stress (Fig. 2A). The variation across PC2 appeared to separate out the transcriptional response to the two warmest acclimation temperatures, 18 and 21 °C. The top categories loading on PC2 in the gill included thermal cellular stress and growth, in opposite directions (Fig. 2A).

**Figure 2.**
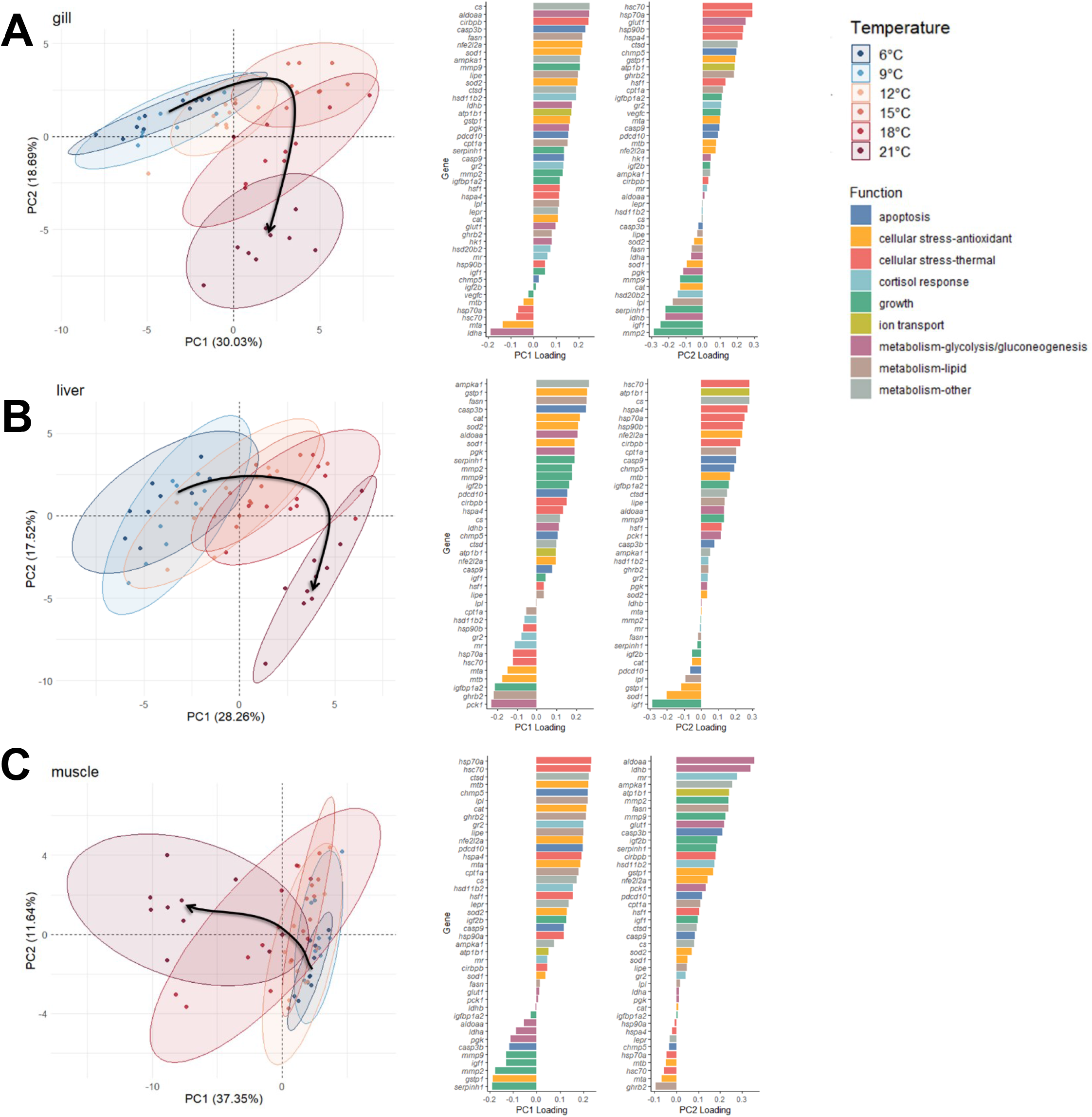
Principal component analysis (PCA) of transcript abundances and their loadings in the gill (A), liver (B), and muscle (C) of bull trout (*Salvelinus confluentus*) acclimated to temperatures ranging from 6 to 21 °C for three weeks. In PCA plots, points represent individual fish (*n* = 9-10), ellipses represent 95% confidence ellipses for each temperature group, and arrows represent the general trend as acclimation temperature increases. The variance in the data explained by each principal component is indicated in brackets. Loadings are plotted for each transcript, describing the contribution of that transcript to the indicated principal component. Transcripts are categorized according to functional role (see Table 1 for full gene names and details).

The liver analysis included 39 target genes. When assessed together by PCA, 28.3% of the variation in the data was explained by PC1 while 17.5% was explained by PC2 (Fig. 2B). Grouping the data by acclimation temperature showed a spread across PC1, with higher temperatures having higher scores. The largest loadings on PC1 in the liver were transcripts involved in the antioxidant response, as well as metabolism (Fig. 2B). Similar to the pattern observed in the gill, the warmest acclimation temperature was separated from all other groups across PC2. As in the gill, the largest loadings on PC2 in the liver included transcripts involved in thermal cellular stress, as well as *atp1b1*, *cs*, and *igf1*.

In the muscle, 42 target genes were analysed. The PCA for the muscle showed very little separation across most acclimation temperatures (6 through 15 °C) while 18 and 21 °C groups were more distinct, primarily having lower scores across PC1 (Fig. 2C). Accordingly, PC1 explained most of the variation in the data (37.4%), while PC2 explained just 11.6%. In the muscle, the highest loadings on PC1 (Fig. 2C) included thermal cellular stress (positive) and growth (negative), like the pattern observed for PC2 in the gill. Additionally, PC1 had strong positive loadings associated with the antioxidant response, and lipid metabolism.

Significant changes in transcript abundance due to increasing acclimation temperature were determined and compiled in figure 3. For simplicity, the changes listed are expressed as the first significant change with increasing acclimation temperature (or decreasing for 6 °C) relative to the 9 °C group, which is closest to their original holding temperature. The supplementary information includes all fold-changes and *p*-values (Tables S1, S2, and S3) and plots (Figures S1, S2, and S3) for gill, liver, and muscle, respectively. This summary demonstrates that the gill experienced the fewest, while the liver had the most transcriptional changes in response to changes in acclimation temperature, including 3 instances of differential regulation (*hsp70a*, *igf1*, *fasn*). Additionally, hepatic transcripts showed at least 3 significantly changed genes at each acclimation temperature relative to 9 °C, relative to other tissues where most changes were observed due to acclimation at 21 °C, particularly in the muscle.

**Figure 3.**
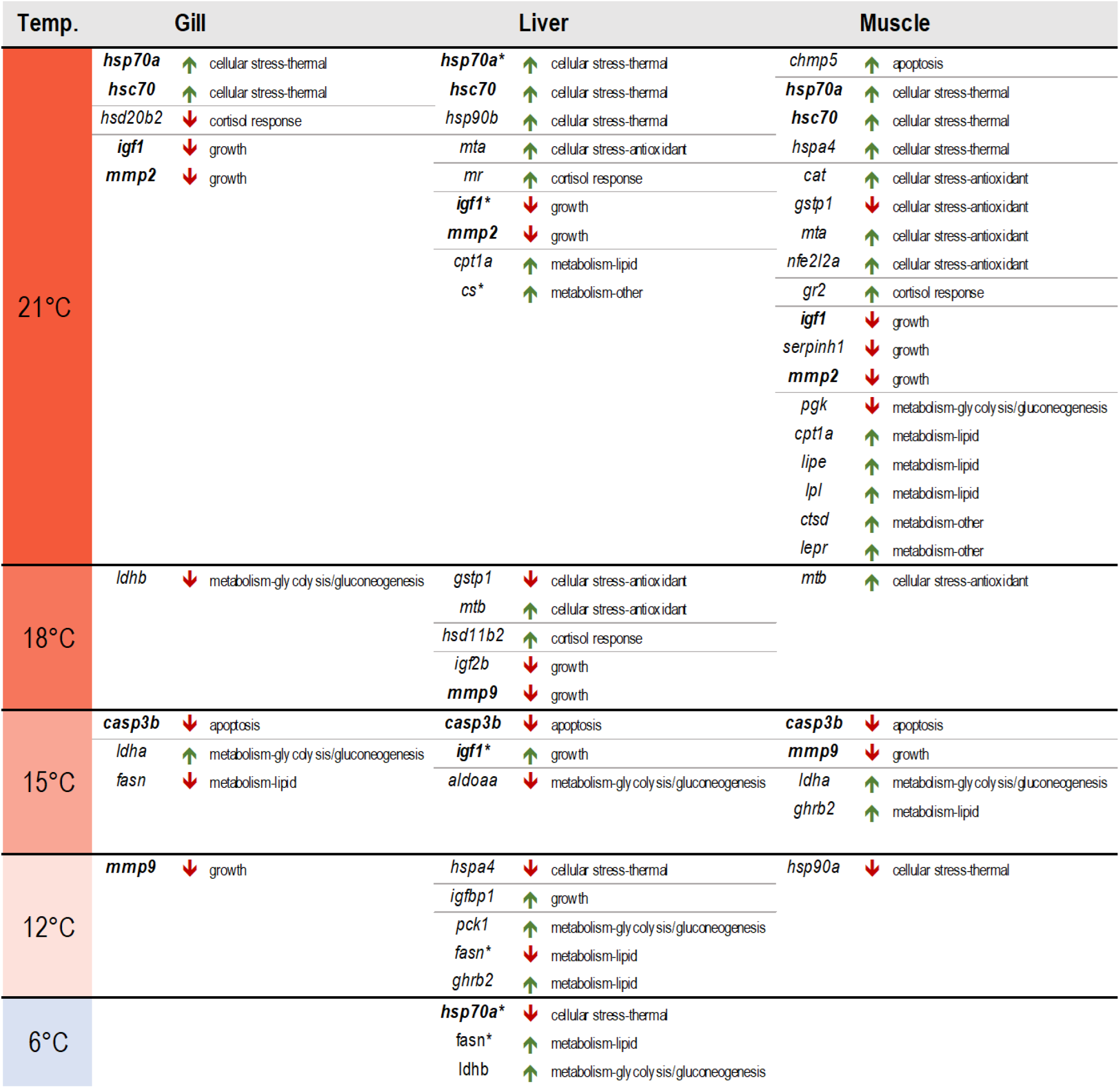
Temperature thresholds for individual transcriptional changes relative to 9 °C by tissue type in bull trout (*Salvelinus confluentus*) acclimated to temperatures ranging from 6 to 21 °C for three weeks. Transcripts are sorted into the acclimation temperature at which they first change significantly relative to 9 °C. Direction of change is indicated by arrows, and functional categories are included according to Table 1. Gene names in bold font indicate significantly changed transcripts common to all 3 tissues, though not necessarily at the same temperature (*casp3b, hsp70a, hsc70, igf1, mmp2, mmp9*). Asterisks indicate transcripts differentially regulated relative to 9 °C and are therefore shown twice (liver *hsp70a, fasn, igf1*).

Several transcripts displayed significant changes in every tissue, including transcripts involved in the thermal stress response (*hsp70a*, *hsc70*), and growth (*igf1*, *mmp2*, *mmp9*). For *hsp70a* and *hsc70*, both were significantly upregulated in every tissue after 3 weeks acclimation at 21 °C (Fig. 4). In the liver, fish acclimated to 6 °C had lower *hsp70a* levels relative to the 9 °C group. Three transcripts involved in growth and development were also significantly changed in all three tissues tested. The matrix metalloproteinases (*mmp*) and insulin-like growth factor 1 (*igf1*) were generally downregulated relative to 9 °C with increasing acclimation temperature, mostly at 21 °C, although *mmp9* downregulation occurred at 12 °C in the gill, 15 °C in the muscle and 18 °C in the liver (Fig. 4). In the liver, *igf1* was differentially regulated, showing a significant upregulation at 15 °C in addition to being downregulated at 21 °C (Fig. 4).

**Figure 4.**
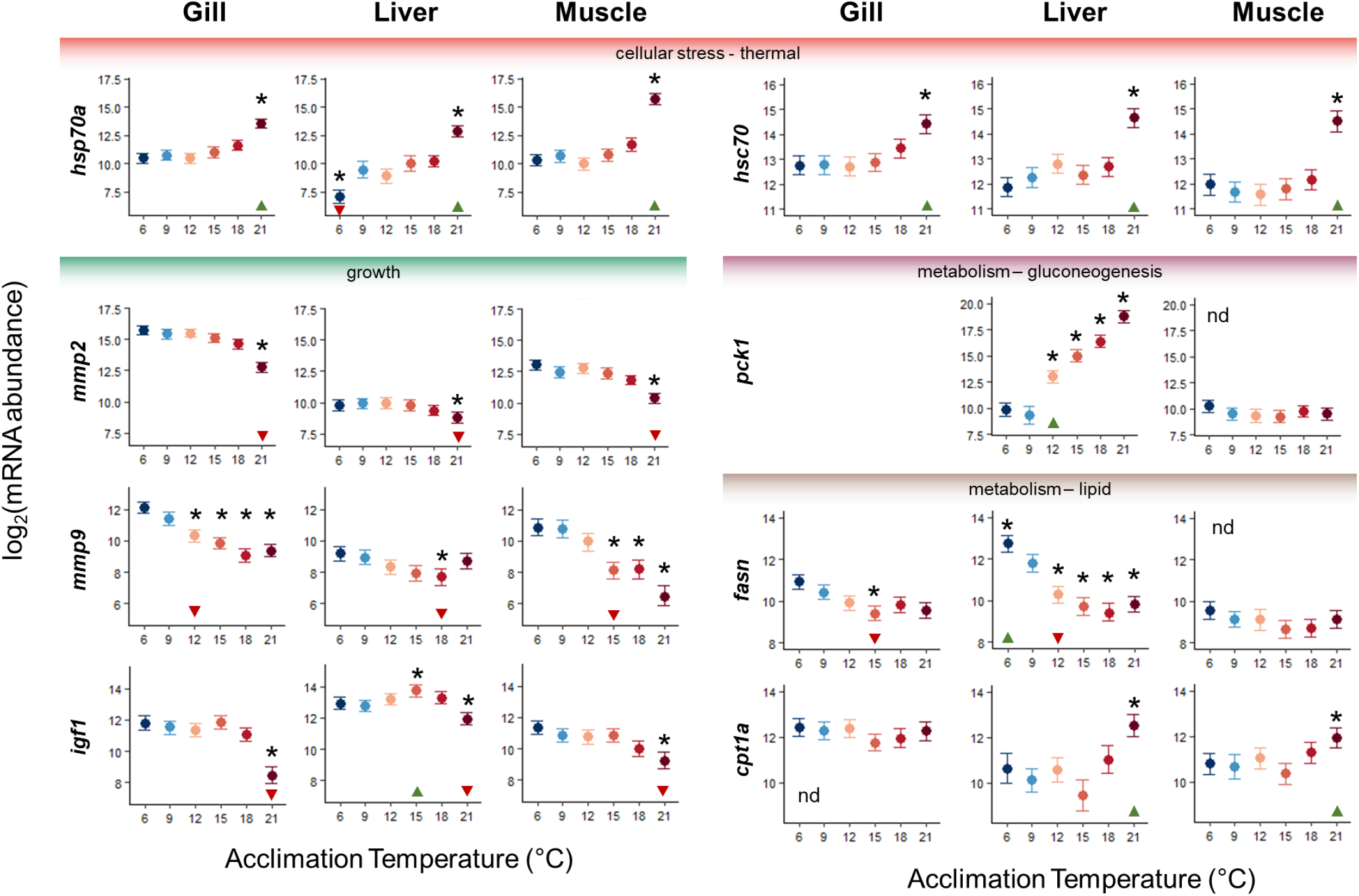
Selected transcript abundance changes across acclimation temperature and tissue type in bull trout (*Salvelinus confluentus*) acclimated to temperatures ranging from 6 to 21 °C for three weeks. Genes pertain to thermal cellular stress (*hsp70a, hsc70*), growth (*igf1, mmp2, mmp9*), and metabolism (*pck1, fasn, cpt1a*) – see Table 1 for gene details. Data are presented as log2(abundance), with points representing means and error bars representing 95% credible intervals. Asterisks indicate significant differences from 9 °C, and triangles along x-axes indicate threshold temperatures listed in figure 3.

In addition to these global results showing similar changes in all tissues, there were also several tissue-specific responses, represented by non-bolded genes in figure 3 with selected genes (*pck1, fasn, cpt1a*) shown in more detail in figure 4. Hepatic phosphoenolpyruvate carboxykinase (*pck1*) expression was perhaps one of the strongest effects in the study, with a significant stepwise increase in abundance with increasing acclimation temperature (Fig. 4). In contrast, *pck1* in the muscle did not change, and abundance in the gill was too low for reliable analysis. Fatty acid synthase (*fasn*), involved in lipid synthesis, was also differentially expressed across tissues, downregulated in the gill at 15 °C and at all temperatures ≥ 12 °C in the liver, but no changes were observed in the muscle (Fig. 4). Additionally, hepatic *fasn* was significantly higher after acclimation to 6 °C, one of the few genes that changed at this cooler temperature.

While *fasn* did not change in muscle tissue, three other transcripts involved in lipid breakdown did, including carnitine palmitoyl transferase (*cpt1a*) which was upregulated in muscle (and liver) as a result of acclimation to 21 °C (Fig. 4).

## 4 Discussion

The bull trout in this study demonstrated substantial differences in their responses to a range of ecologically-relevant water temperatures. While the capacity for acclimation based on *CT_max_* and whole-body performance metrics (growth, mortality) clearly identified 21 °C as a physiologically stressful temperature for bull trout, effects were also apparent at lower temperatures, particularly at a transcriptional level. The transcriptional response profiles to different acclimation temperatures included genes that changed consistently in all 3 tissues, with significant changes in these transcripts mostly (but not exclusively) occurring at 21 °C, when thermal stress was evident. In contrast, several transcripts had clear tissue-specific patterns which speaks to their functional role during acclimation to a given temperature. For instance, the liver appears to be regulating metabolic pathways at colder acclimation temperatures (6 °C) as well as in response to warming acclimation temperatures (12-15 °C), while transcriptional changes in the muscle mostly occur alongside indicators of thermal stress, around 21 °C. These lower thresholds apparent in the liver likely represent adaptive responses to thermal stress (i.e. metabolic adjustments allowing the animal to redirect energy based on their environmental temperature) while the 21 °C changes in the muscle represent sublethal thermal thresholds.

Bull trout demonstrated a loss of the capacity for improvement of thermal tolerance via acclimation at 21 °C, suggesting there is an important sublethal thermal threshold between 18 and 21 °C. Beyond 18 °C, the biological processes affected by acclimation to elevated temperatures are no longer capable of increasing *CT_max_*. The positive impact of increasing acclimation temperature on acute thermal performance is well known in fishes; however, this can be influenced by many variables with respect to the fish (species, population, life stage, genetic background), and acclimation regime (Schulte et al. 2011; McKenzie et al. 2021; Ern et al. 2023). For example, a recent study on brook trout (*Salvelinus fontinalis*) found that *CT_max_* increased with acclimation temperature until reaching a plateau of ∼31.8 °C at acclimation temperatures beyond 20 °C (Morrison et al. 2020). A previous study in age-0 bull trout also found improved thermal tolerance at higher acclimation temperatures with a mean *CT_max_* of 26.4 to 28.9 °C after two weeks of acclimation to 8, 12, 16, and 20 °C (Selong et al. 2001). In that study, a plateau was not obvious and *CT_max_* values were somewhat lower than in the current study.

Transcript abundance data suggest a broad shift in transcriptional regulation in all tissues measured at 21 °C. The PCAs for gill and liver demonstrate that this higher-temperature shift is driven by changes in genes involved in thermal stress and growth, as these had the highest loadings on PC2. The shift in PC1 in gill and liver suggest a broader thermal shift in transcription that encompasses lower acclimation temperatures as well, driven by changes in metabolic and antioxidant genes. In contrast, this 21 °C shift is the primary trend (PC1) observed in the muscle, showing separation from the remaining groups that cluster together, with 18 °C being intermediate. This distinct transcriptional shift in all tissues in the 21 °C group, combined with having the lowest growth and survival rates, suggests that at this temperature, bull trout are experiencing chronic thermal stress. In support of a chronic thermal stress response, the heat shock proteins loaded strongly onto PC2 (and to a lesser extent onto PC1 for muscle), suggesting that these fish are chronically and widely activating this cellular stress response to cope with this elevated temperature. Heat shock proteins are chaperones that aid in the folding of polypeptides as they are synthesized, and mediate the repair and degradation of proteins that are damaged, by heat denaturation for instance (Basu et al. 2002). Several studies have observed changes in the heat shock proteins due to increasing acclimation temperature. For instance, 5 months of acclimation to 18 °C increased basal *hsp70* (but not *hsc70*) in gill, liver, and white muscle of juvenile lake whitefish (*Coregonus clupeaformis*) and warm-acclimated (17 °C) rainbow trout (*Oncorhynchus mykiss*) had higher basal *hsp70* mRNA in the blood (Currie et al. 2000). The constitutive *hsc70* has also been shown to have elevated expression with increasing acclimation temperature in brook trout, another char species (Stitt et al. 2014; Mackey et al. 2021), while (Fangue et al. 2006) showed higher expression in a warmer southern population of killifish compared to their northern counterparts. Generally, the impact of acclimation temperature on expression patterns of the heat shock protein family members remains complex, depending on species, tissue, and thermal history. However, the robust upregulation observed at 21 °C in the current study points to a widespread cellular thermal stress response in bull trout.

Increasing acclimation temperature also appears to be suppressing growth. Relative growth rates of bull trout peaked at a temperature (10.9 °C) within their original rearing temperature (10 °C ± 1 °C), and dropped most dramatically at 21 °C, to negative growth. Further, insulin-like growth factor 1 (*igf1*) loaded strongly in an opposing direction to the heat shock proteins in all tissues and is significantly downregulated at 21 °C in all tissues. The primary source of circulating IGF-I is the liver, where this hormone is produced in response to growth hormone from the pituitary and acts to stimulate somatic growth in fish. IGF-I is also produced outside the liver, where its effects are thought to be more local, i.e. autocrine/paracrine signalling (Reinecke 2010). IGF binding proteins (*igfbp*) bind circulating IGF to reduce its growth effects, often while catabolism is enhanced (Reinecke 2010). In the current study, hepatic *igfbp1* displayed a robust pattern of increase with acclimation temperature, which may further reduce *igf1*-mediated growth signalling. Temperature is known to have an impact on growth signalling, with fish reared at higher temperatures within their tolerance zone generally having elevated *igf1* levels (Reinecke 2010). Indeed, bull trout did have elevated hepatic *igf1* at 15 °C. However, prolonged thermal elevation beyond this range led to reduced growth and food intake. For example, and in agreement with current findings, bull trout held at 20 °C had markedly reduced growth and fed very little (Selong et al. 2001). In another study, Atlantic salmon (*Salmo salar*) exposed to elevated temperatures for 15 and 45 days had reduced muscle *igf* mRNA and increased liver *igfbp* (Hevrøy et al. 2012), both of which were also observed in the current study.

Another set of genes related to growth that displayed downregulation as acclimation temperature increased in all tissues are the matrix metalloproteinases (*mmp*). These are collagenases involved in tissue remodelling via extracellular matrix degradation either during normal developmental processes or in disease states (Visse and Nagase 2003). Zebrafish acclimated to 34 °C had reduced growth and reduced transcript abundance of muscle collagenases, including *mmp2* and *mmp9* (Lin et al. 2022). Similarly, both were downregulated in bull trout muscle, as well as gill and liver, as acclimation temperature increased in the current study. In bull trout muscle and gill, collagen-related processes loaded strongly opposite the heat shock response (onto PC1 or PC2, respectively), implicating a reduced capacity for tissue remodelling at elevated temperatures. Overall, growth processes appear to be most impacted in bull trout exposed to 21 °C, based on both relative growth rate and transcript data. However, some changes were observed as early as 12 °C (*igfbp1* upregulation in the liver) or 15 °C (liver *igf1* and muscle *mmp9*), potentially due to metabolic adjustments, suggesting that impacts on growth are detectable at lower acclimation temperatures. Among the wide range of factors regulating growth and development, these processes are most deeply intertwined with metabolic status. In line with this, other transcriptional changes with respect to metabolism were also observed, and tended to be more sensitive indicators of increased temperature.

In contrast to the liver, metabolic changes in the muscle are mostly associated with a more critical sublethal threshold at 21 °C. Muscle PC1 captures most of the variation in the data, and while heat shock and growth genes are most heavily loaded, there are several genes loading strongly onto PC1 involved in oxidative stress and lipid metabolism. These changes generally suggest increases in lipid (*cpt1a*, *lipe*, *lpl*, *lepr*) and protein (*ctsd*) breakdown in the muscle, changes which are balanced by reductions in growth indicators. Hepatic *cpt1a* was also upregulated at 21 °C, suggesting enhanced use of lipid as metabolic fuel in this tissue. Fatty acid synthase (*fasn*), which mediates lipogenesis, tended to be lower at warmer acclimation temperatures in gill and liver and was higher in the liver at 6 °C, one of the few changes associated with cold acclimation. Together, this suggests that thermal stress is having an impact on lipid metabolism in all tissues. In support of this, brook trout acclimated to higher temperatures had less visceral adipose tissue (Morrison et al. 2020). These metabolic changes may be exacerbated by reduced feeding, and while food intake was not specifically tracked in this study, bull trout tend to reduce their feed intake at temperatures warmer than 16 °C (Selong et al. 2001; Mesa et al. 2013). The changes in lipid metabolism observed at both 6 and 21 °C suggest that fish are more sensitive at these temperature extremes, and may help explain why bull trout are not often observed in streams below 6 °C, despite their accessibility. The transcriptional data suggest that fish at 21 °C may be directing lipid and protein stores toward fueling elevated metabolic rates and upregulated cellular stress responses at the expense of growth, which is lowest at 21 °C.

The changes observed in agitation temperature during this study were less straightforward than the *CT_max_*. Agitation temperature did not increase with increasing acclimation temperature in the current study. While an increase in agitation temperature has been observed with increasing acclimation temperature in a previous study on pugnose shiner, *Notropis anogenus* (McDonnell et al. 2021), agitation temperature can also be stable across different acclimation temperatures (despite changes in *CT_max_*); as observed in both an African cichlid, *Pseudocrenilabrus multicolor* (McDonnell and Chapman 2015) and Athabasca rainbow trout along a temperature gradient (Hnytka 2023). Similarly, agitation temperature did not vary while *CT_max_* did (with summer temperature), across different populations of wild brook trout (Wells et al. 2016). Based on the handful of studies to date looking at agitation temperature, this metric of thermal tolerance seems to be less environmentally plastic than *CT_max_*. While the limited capacity for acclimation is an important sublethal threshold, *CT_max_* itself is an important proxy of the lethal thermal limits. Bull trout in the current study recovered from the *CT_max_* protocol, suggesting that the true lethal thermal limit was not exceeded. The *CT_max_* plateau observed in this study (29.3 ± 0.1 °C) may represent a ‘hard ceiling’ of acute upper thermal tolerance for bull trout, at least through acclimation at this life stage in the current population.

Adaptation to increasing temperatures could theoretically raise this ceiling (Ern et al. 2023).

Altogether, bull trout showed evidence of thermal acclimation and thermal stress at both the whole-organism and molecular level across an acclimation gradient spanning from 6 to 21 °C. By all metrics, bull trout did not perform well at 21 °C: their *CT_max_* is at its plastic limit, growth and survival are poor, and there is a large transcriptional response across all tissues tested in terms of cellular stress and growth. The response to an acclimation temperature of 18 °C does not appear as severe, with *CT_max_* being higher than the previous acclimation temperature (15 °C), a milder effect on growth but not survival, and fewer signs of stress based on transcript abundance. Together, this suggests that 18 °C is an important sublethal thermal threshold, beyond which fish performance suffers. With the exception of agitation temperature at 6 °C, the fish performance endpoints we measured were consistent below 18 °C, but PCA of transcriptional data points to metabolic changes occurring in response to changes in acclimation temperature, generally suggesting a shift to catabolism, starting as early as 12 °C in the liver.

Several transcripts relating to cellular stress and growth were altered at 21 °C regardless of tissue type, providing potentially useful ‘global’ markers of thermal stress, while tissue-specific effects highlighted different aspects of thermal acclimation, particularly in liver and muscle.

Wild bull trout are rarely found at temperatures above 11 °C, and though this likely varies by population, this observation corresponds well to the data in the current study, with detrimental impacts occurring mainly beyond 15 °C (Jones et al. 2014; Mochnacz et al. 2023). While bull trout have been observed (temporarily) at 20.5 °C, it is likely not ideal based on growth and survival in the current and previous studies (Selong et al. 2001), and the transcriptional response at 21 °C in the current study. However, the results of this study are limited to YOY bull trout from one wild population, reared in a controlled environment. Thermal tolerance can differ across genetic populations and life stages, with embryos and spawning adults tending to be more sensitive (McKenzie et al. 2021). Additionally, fish in this study were held at relatively constant temperatures whereas wild fish may experience thermal variation, potentially exacerbated by climate change (e.g. increased peak temperatures), which may impact performance (Morash et al. 2021). Therefore, exploration of these factors would further clarify and refine our knowledge of the impacts of climate warming on bull trout. Despite these caveats, our study provides evidence of when water temperatures begin to become harmful to bull trout, which can be used to inform management strategies for wild populations.

## Acknowledgments

Thank you to Kerry Wautier, Myles Panciera and Brennan Romaniuk for help with fish holding and husbandry and Sarah Hnytka for helping with the *CT_max_* experiments.

## Author contributions

- CB: Conceptualization; Methodology; Validation; Formal Analysis; Investigation; Writing-Original Draft; Visualization; Writing - Review & Editing
- TCD: Conceptualization; Methodology; Validation; Investigation; Writing – Original Draft; Writing - Review & Editing
- AJC: Conceptualization; Methodology; Investigation; Writing - Review & Editing
- MA: Investigation; Writing - Review & Editing
- SSI: Methodology; Writing - Review & Editing
- NJM: Conceptualization; Methodology; Resources; Supervision; Project Administration; Funding Acquisition; Writing - Review & Editing
- DDH: Funding Acquisition; Methodology; Writing - Review & Editing
- KMJ: Conceptualization; Methodology; Resources; Supervision; Project Administration; Funding Acquisition; Writing - Review & Editing

## Data Availability

Data are available in the supplementary information (Tables S4 and S5), and in the GitHub repository with associated R scripts used for analysis at https://github.com/c3best/Bull_Trout_Transcript_Analysis.

## Graphical Abstract / Figure X

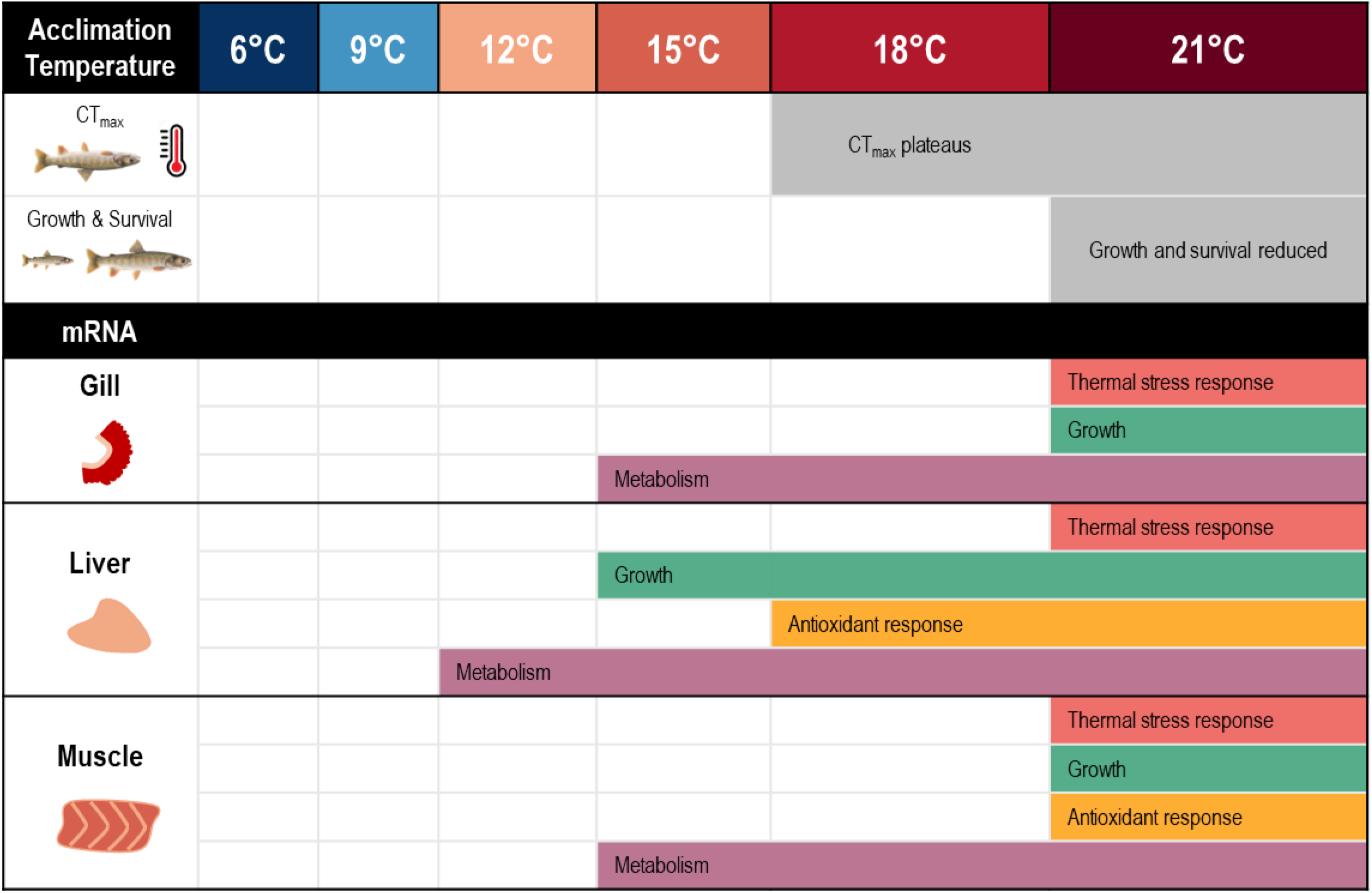

